# The Network Analysis Profiler (NAP v2.0): A web tool for visual topological comparison between multiple networks

**DOI:** 10.1101/2020.11.14.382580

**Authors:** Mikaela Koutrouli, Theodosios Theodosiou, Ioannis Iliopoulos, Georgios A. Pavlopoulos

## Abstract

In this article we present the Network Analysis Profiler (NAP v2.0), a web tool to directly compare the topological features of multiple networks simultaneously. NAP is written in R and Shiny and currently offers both 2D and 3D network visualization as well as simultaneous visual comparisons of node- and edge-based topological features both as bar charts or as a scatterplot matrix. NAP is fully interactive and users can easily export and visualize the intersection between any pair of networks using Venn diagrams or a 2D and a 3D multi-layer graph-based visualization. NAP supports weighted, unweighted, directed, undirected and bipartite graphs and is available at: http://bib.fleming.gr:3838/NAP/. Its code can be found at: https://github.com/PavlopoulosLab/NAP

## 1 Introduction

Networks are key representations which can capture the associations and interactions between any kind of bioentity such as genes, proteins, diseases, drugs, small molecules and others (1–6). Gene co-expression networks, gene regulatory networks, protein-protein interaction networks (PPIs), signal transduction networks, metabolic networks, gene-disease networks, sequence similarity networks, phylogenetic networks, ecological networks, epidemiological networks, drug-disease networks, disease-symptom networks, literature co-occurrence networks, food webs, semantic and knowledge networks are the most widely known network types in the biomedical and biomedical-related fields (6). However, not all networks are the same in terms of structure and come with certain topological features. Erdos–Rényi (7) networks for example are random graphs with no specific structure, Watts-Strogatz (8) networks are random graphs with small communities and Barabási–Albert (9) networks are random scale-free networks whose degree distribution follows a power law. While basic topological network analysis is offered by widely used network visualization applications (3) such as the Cytoscape (10) and Gephi (11), in this article, we present NAP v2.0, a complementary web-based tool designed to fill certain gaps and aid non-experts in not only analyzing the topological features of a network but to visually perform direct comparisons across multiple network in an easy and user-friendly way.

## 2 The application

In its current version, NAP (12) supports weighted, unweighted, directed, undirected simple and bipartite networks. It is implemented in R and Shiny and most of its backend calculations are based on the *igraph* library (13). It accepts as input a tab-delimited file in which the first two columns indicate the connections between the nodes and the third column the weight between these edges. Users have the option to upload as many networks as they like, name them accordingly and process them simultaneously. NAP v2.0 has four main functions: *i)* Basic Visualization, *ii)* Topological analysis, *iii)* Node/Edge ranking and *iv)* Intersection network hosting the common vertices and edges between two selected networks.

### 2.1 Basic 2D/3D visualization

Once a network has been uploaded and named, it is visualized with the use of *visNetwork* library. *VisNetwork* offers a fully interactive visualization as it allows network zooming, dragging and panning. Nodes can be selected and placed anywhere on the plane, whereas the first neighbors of any node can be highlighted upon selection. This network view can show one network at a time and is automatically updated when a different network is selected. In this view, NAP supports the following *igraph* layouts (14):

- Fruchterman-Reingold (15): It places nodes on the plane using a force-directed layout.
- *Random*: This function places the vertices on a 2D plane uniformly using random coordinates.
- *Circle*: It places vertices on a circle, ordered by name.
- Kamada-Kawai (16): This layout places the vertices on a 2D plane by simulating a physical model of springs.
- Reingold-Tilford (17): This is a tree-like layout and is suitable for trees, ontologies and hierarchies.
- LGL (18): A force directed layout suitable for larger graphs.
- *Grid*: This layout places vertices on a rectangular 2D grid.
- *Sphere*: This layout places vertices on a rectangular 3D-like sphere.

In addition to the 2D visualization, NAP offers a fully interactive 3D network visualization using a force-directed layout. Users can zoom-in and out and interactively drag and drop a node or the whole network and place it anywhere in space. This visualization is based on the *D3*.*js* library and is sufficient for larger graphs, especially in cases where the 2D view becomes overcrowded. An example of a Yeast PPI (19) is shown in Figure 1.

**Figure 1:**
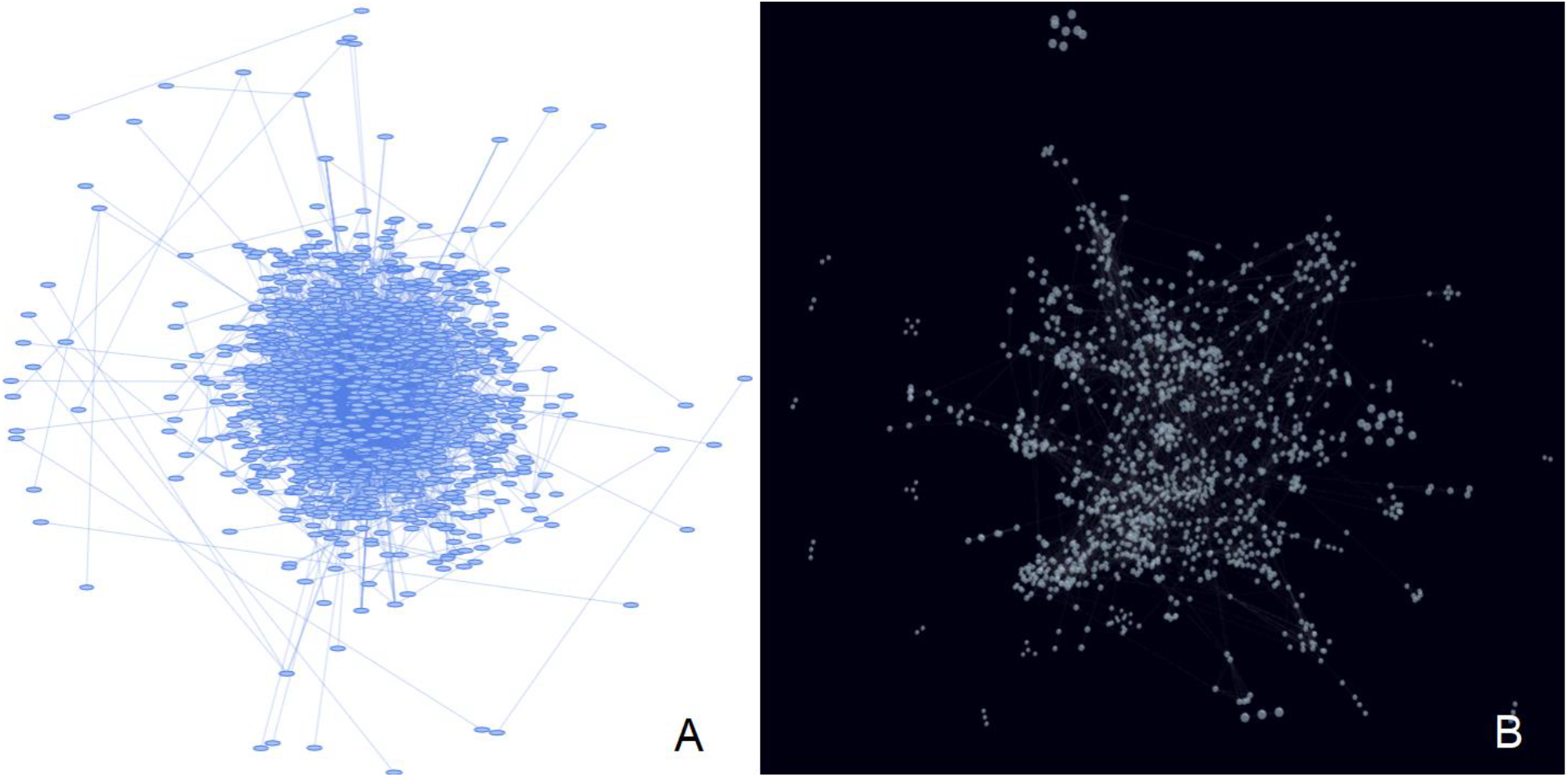
NAP’s basic visualization. *A*) 2D visualization of a Yeast PPI (19) using the Kamada-Kawai layout. *B)* The same network visualized in 3D.

### 2.2 The topological features

In its current version, NAP supports the following *igraph*-based topological features:

- *Number of nodes*: Shows the number of vertices in the network.
- *Diameter*: Shows the length of the longest geodesic. The diameter is calculated by using a breadth-first search like method. The graph-theoretic or geodesic distance between two points is defined as the length of the shortest path between them.
- *Radius*: The eccentricity of a vertex is its shortest path distance from the farthest other node in the graph. The smallest eccentricity in a graph is called its *radius*.
- *Density*: The density of a graph is the ratio of the number of edges divided by the number of possible edges.
- *Average path length*: The average number of steps needed to go from a node to any other.
- *Clustering Coefficient*: A metric to show if the network has the tendency to form clusters.
- *Modularity*: This function calculates how modular is a given division of a graph into subgraphs.
- *Number of self-loops*: How many nodes are connected to themselves.
- *Average Eccentricity*: The eccentricity of a vertex is its shortest path distance from the farthest other node in the graph.
- *Average Eigenvector Centrality*: It shows the influence of a node in a network.
- *Assortativity degree*: The assortativity coefficient is positive if similar vertices (based on some external property) tend to connect to each or negative otherwise.
- *Directed acyclic graph*: It shows if a graph has cycles or not.
- *Average number of Neighbors*: How many neighbors each node of the network has on average.
- *Centralization betweenness*: It is an indicator of a node’s centrality in a network. It is equal to the number of shortest paths from all vertices to all others that pass through that node. Betweenness centrality quantifies the number of times a node acts as a bridge along the shortest path between two other nodes.
- *Centralization closeness*: It measures the speed with which randomly walking messages reach a vertex from elsewhere in the graph.
- *Centralization degree*: It is defined as the number of links incident upon a node.
- *Graph mincut:* Calculates the minimum *st*-cut between two vertices in a graph. The minimum *st*-cut between source and target is the minimum total weight of edges needed to remove to eliminate all paths from source to target.
- *Motifs-3:* Searches a graph for motifs of size 3 (6).
- *Motifs-4*: Searches a graph for motifs of size 4 (6).

While users can select and visualize each topological measure in a numeric form, one can select several of the uploaded networks and directly compare their topological features in different bar charts. Figure 2 shows an example of a direct comparison between two Yeast PPI networks (19, 20) (generated in 2002 and 2006 respectively) and a random scale-free Albert-Barabasi network consisting of 1000 nodes (generated by NAP’s automatic network generators). Bar charts are fully interactive and are produced with the use of the *plotly* library.

**Figure 2:**
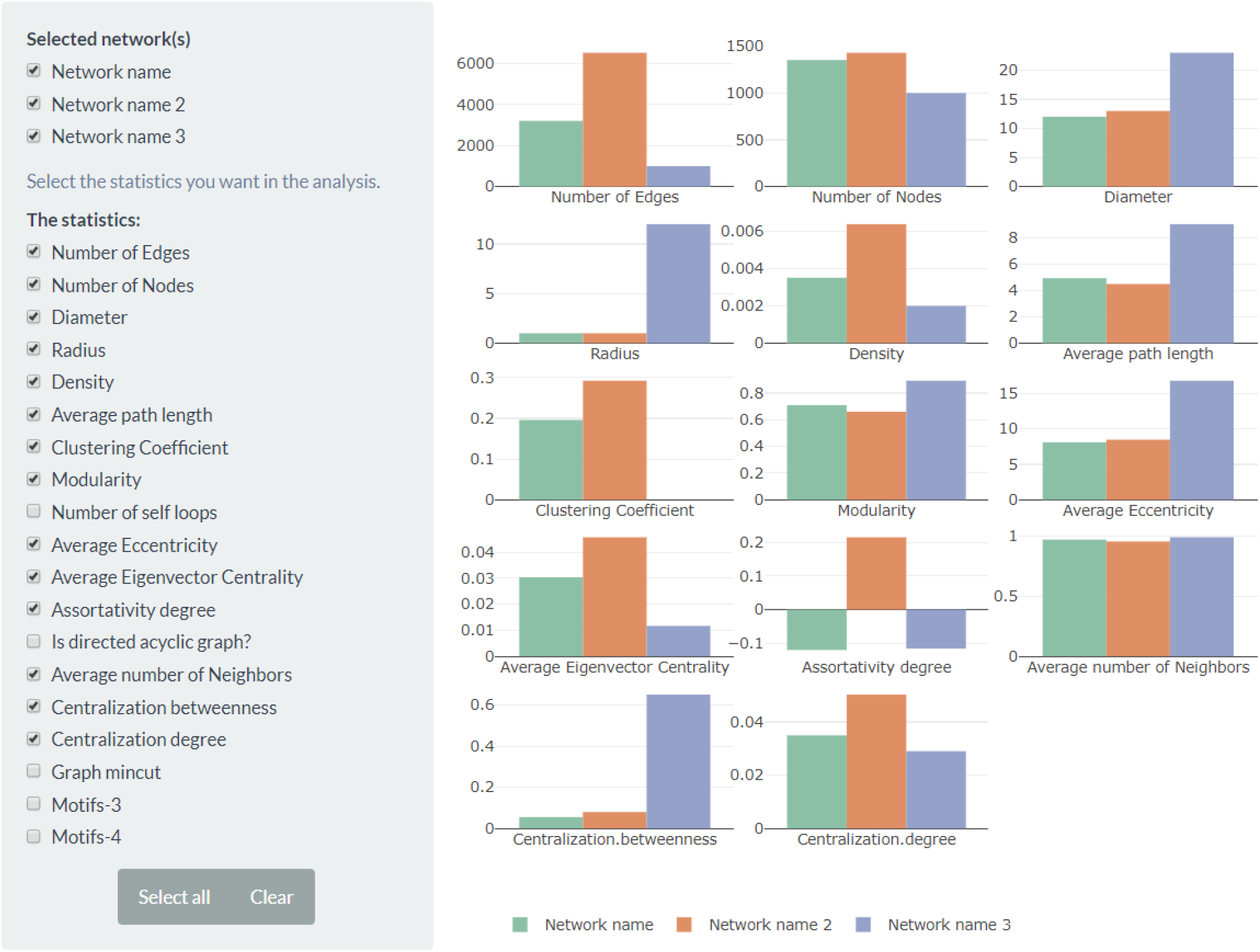
Direct comparison of fourteen topological features across three different networks.

### 2.3 Topological feature pairwise comparison and node/edge ranking

As explained in NAP’s v1.0 article (12), nodes can be ranked by *centralization degree*, *centralization betweenness*, *clustering coefficient*, *page rank*, *eccentricity*, *eigenvector* and *subgraph* centrality whereas edges can be ranked by *betweenness centrality* only. An all-versus-all scatterplot matrix can be generated to show the pairwise correlations between any of the selected topological features (Figure 3). The upper part of the matrix shows the correlation between any pair of features in a numerical form whereas its lower part shows these correlations in a scatterplot. In the case where only one option has been selected, the viewer will generate a chart showing the values of the selected topological feature in a histogram.

**Figure 3:**
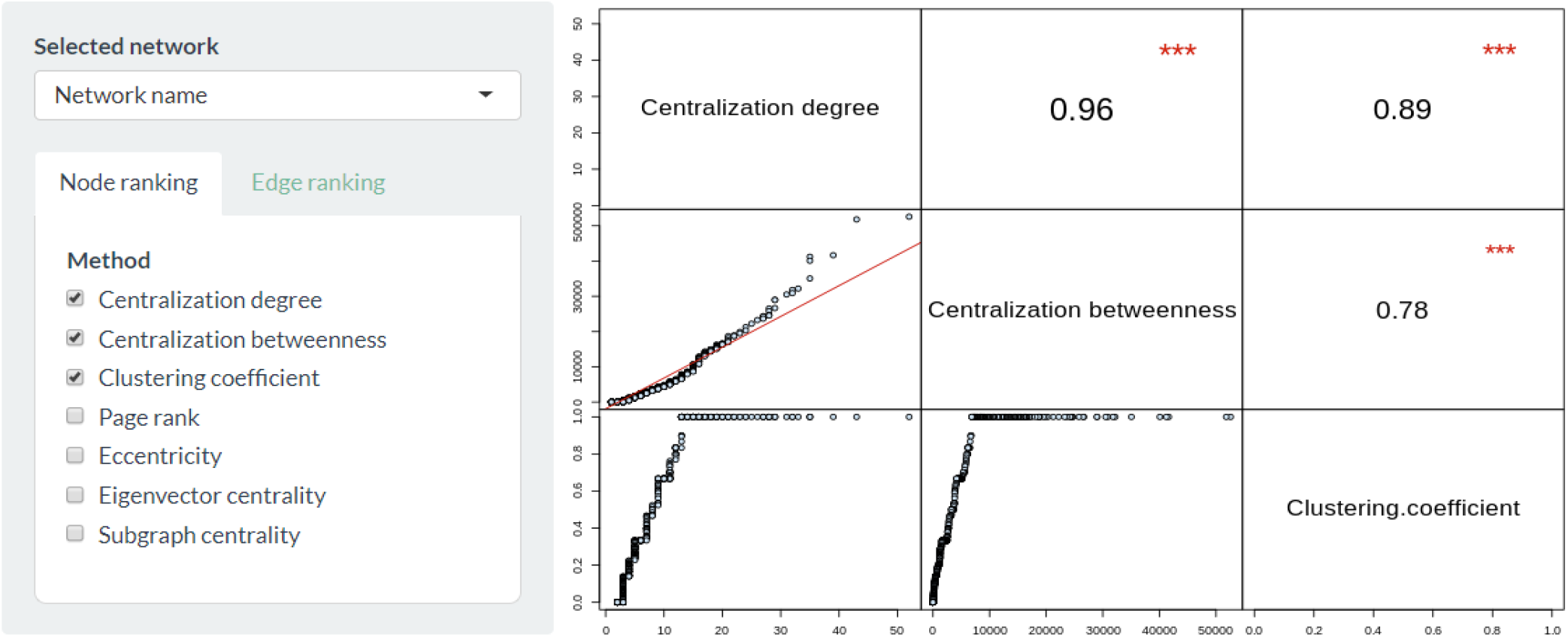
Intra-network pairwise topological feature comparison.

### 2.4 Network intersection

With NAP, users can automatically extract, export and visualize the common edges and nodes between any selected pair of networks. Common node and edge names will be initially reported in interactive tables as text whereas Venn diagrams are used to show the node/edge union and intersection between the two networks. Venn diagrams are generated with the use of *Venndiagrams* library whereas *VisNetwork* library is used to visualize the network’s intersection in an interactive 2D view. In Figure 4, a comparison between two Yeast PPI networks (19, 20), generated in 2002 and 2006 respectively is shown.

**Figure 4:**
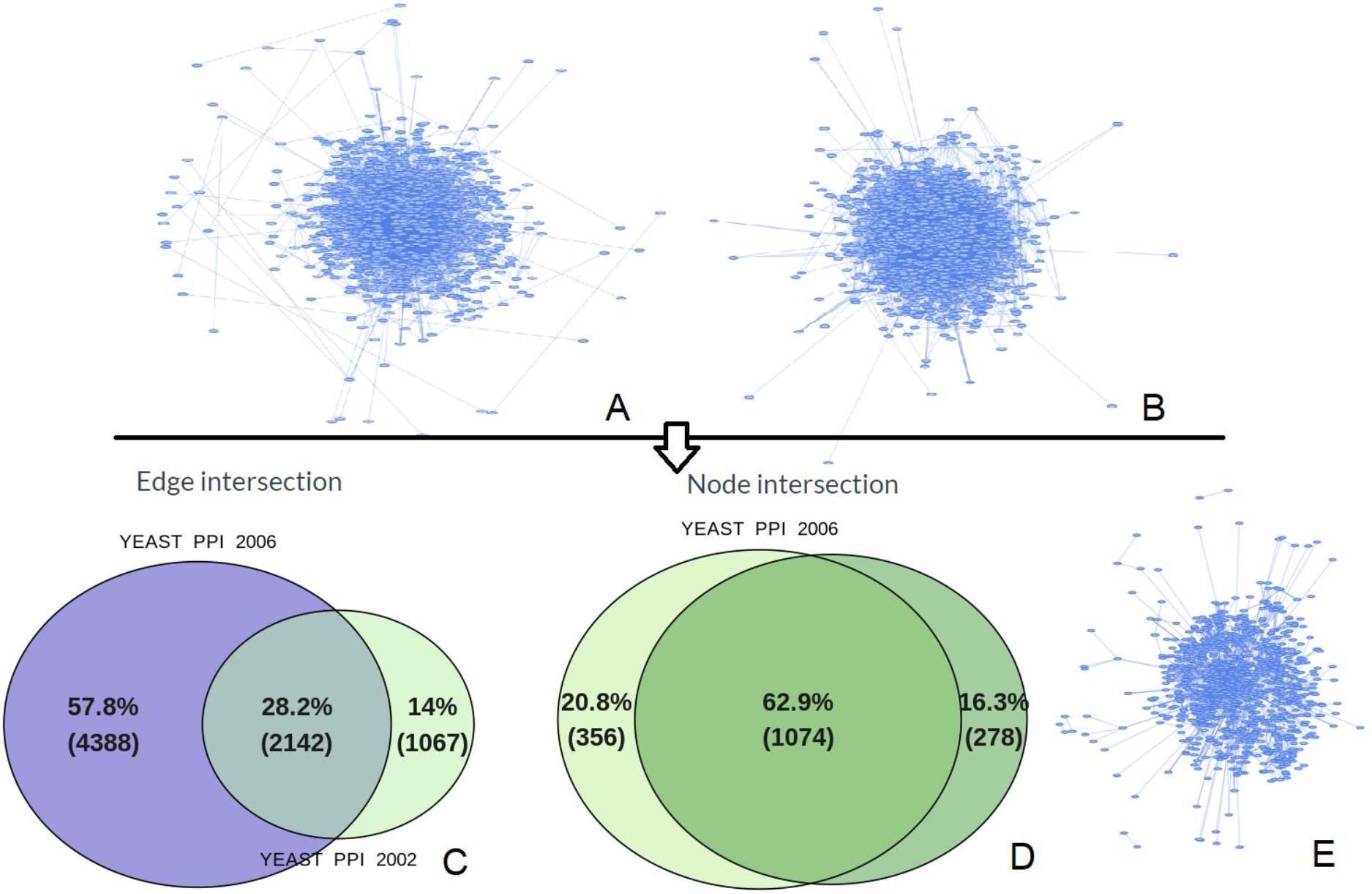
Automatic generation of common edges and nodes between two selected networks. A) A Yeast PPI network generated in 2002 (19). B) A Yeast PPI network generated in 2006 (20). C) 2142 common edges shown in a Venn diagram. D) 1074 common nodes shown in a Venn diagram. E) The generated network consisting of the common edges between the 2002 and 2006 yeast PPI networks.

In addition to the 2D view, NAP gives the option to visualize the common parts between two selected networks using a 3D multi-layer graph implemented in *D3.js*. Nodes of the first network are placed on a layer and are colored blue whereas nodes from the second network are placed on a different layer and are colored red. Nodes which belong to the two different networks but have the same name are considered as common and are colored yellow whereas edges are drawn to connect these nodes across the two layers. Notably, users can use a 3-layer representation to place the common nodes on a third middle layer for a more comprehensive view (not always better).

In order to minimize the crossovers between the lines across layers, a layout can be initially applied on the whole network and nodes can be separated on their two distinct layers upon completion by adjusting their height coordinate. The layouts which are currently supported by NAP for this view are the: *random*, *circular*, *fruchterman-reingold, fruchterman-reingold grid*, *kamada-kawai*, *spring*, and *LGL* force directed algorithm.

The multi-layer 3D graph is fully interactive and users can zoom in/out and drag and rotate each node or the whole network in 3D space for easier exploration. In addition, users can export the network in a text file in order to be processed by more advanced third-party 3D visualizers like, for example, Arena3D (21, 22). The whole concept is schematically shown in Figure 5.

**Figure 5:**
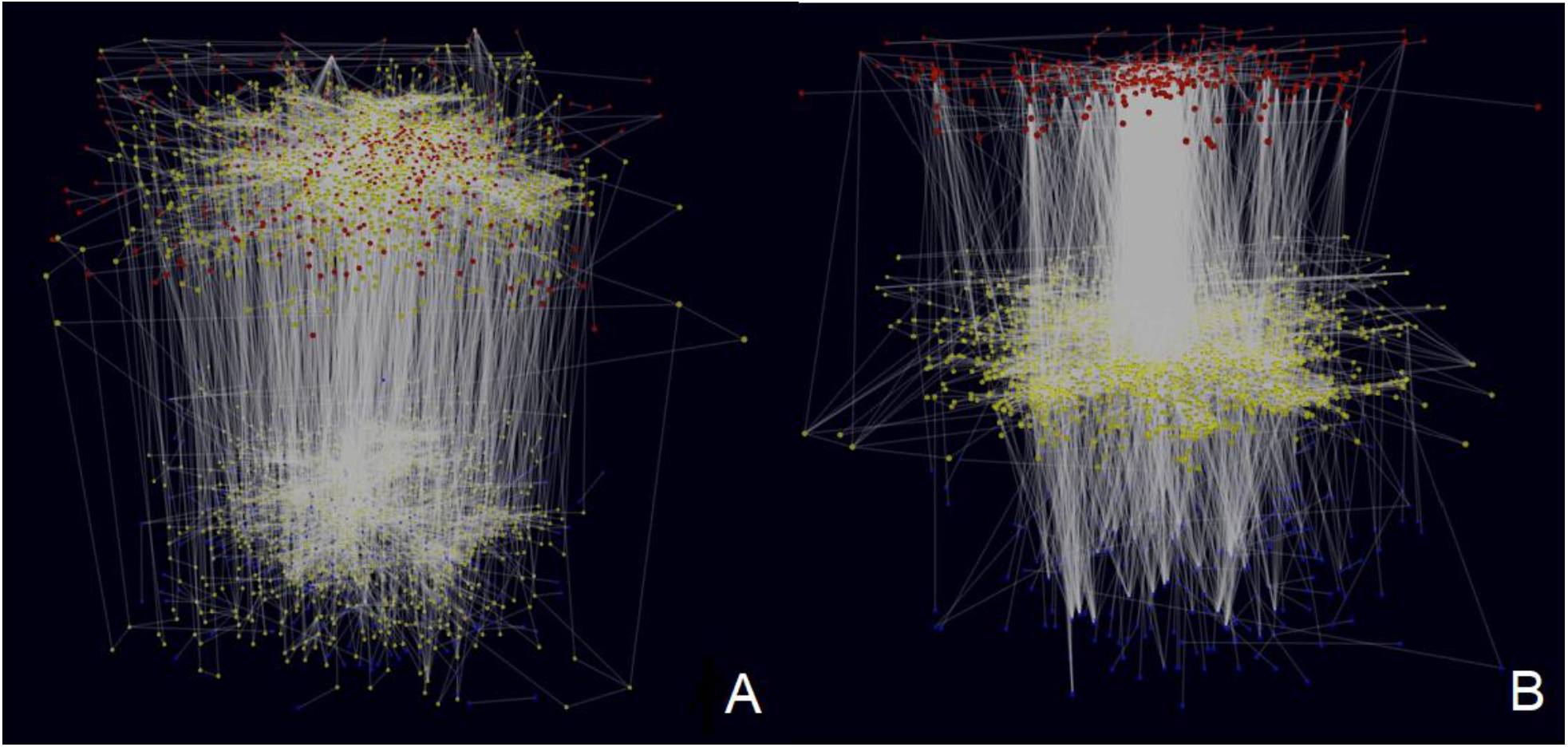
Visualization of common nodes between two networks using a 3D multi-layer graph. A) Two Yeast PPI networks (2002 and 2006 accordingly) are placed in two different layers and their common nodes are marked yellow. B) Alternatively, common nodes are separated and placed on a third middle layer.

## ACKNOWLEDGMENTS

GAP and MK were supported by the Hellenic Foundation for Research and Innovation (H.F.R.I) under the “First Call for H.F.R.I Research Projects to support Faculty members and Researchers and the procurement of high-cost research equipment grant”, GrantID: 1855-BOLOGNA.

## REFERENCES

1. Pavlopoulos,G.A., Secrier,M., Moschopoulos,C.N., Soldatos,T.G., Kossida,S., Aerts,J., Schneider,R. and Bagos,P.G. (2011) Using graph theory to analyze biological networks. BioData Min., 4, 10.

2. Pavlopoulos,G.A., Malliarakis,D., Papanikolaou,N., Theodosiou,T., Enright,A.J. and Iliopoulos,I. (2015) Visualizing genome and systems biology: technologies, tools, implementation techniques and trends, past, present and future. GigaScience, 4, 38.

3. Pavlopoulos,G.A., Wegener,A.-L. and Schneider,R. (2008) A survey of visualization tools for biological network analysis. BioData Min., 1, 12.

4. Pavlopoulos,G.A., Iacucci,E., Iliopoulos,I. and Bagos,P. (2013) Interpreting the Omics ‘era’ Data. In Tsihrintzis,G.A., Virvou,M., Jain,L.C. (eds), Multimedia Services in Intelligent Environments. Springer International Publishing, Heidelberg, Vol. 25, pp. 79–100.

5. Kontou,P.I., Pavlopoulou,A., Dimou,N.L., Pavlopoulos,G.A. and Bagos,P.G. (2016) Network analysis of genes and their association with diseases. Gene, 590, 68–78.

6. Koutrouli,M., Karatzas,E., Paez-Espino,D. and Pavlopoulos,G.A. (2020) A Guide to Conquer the Biological Network Era Using Graph Theory. Front. Bioeng. Biotechnol., 8, 34.

7. Bollobás,B. (2001) Random graphs 2nd ed. Cambridge University Press, Cambridge; New York.

8. Watts,D.J. and Strogatz,S.H. (1998) Collective dynamics of ‘small-world’ networks. Nature, 393, 440–442.

9. Barabási,A.-L. and Albert,R. (1999) Emergence of Scaling in Random Networks. Science, 286, 509–512.

10. Shannon,P., Markiel,A., Ozier,O., Baliga,N.S., Wang,J.T., Ramage,D., Amin,N., Schwikowski,B. and Ideker,T. (2003) Cytoscape: a software environment for integrated models of biomolecular interaction networks. Genome Res., 13, 2498–2504.

11. Bastian,M., Heymann,S. and Jacomy,M. (2009) Gephi: An Open Source Software for Exploring and Manipulating Networks.

12. Theodosiou,T., Efstathiou,G., Papanikolaou,N., Kyrpides,N.C., Bagos,P.G., Iliopoulos,I. and Pavlopoulos,G.A. (2017) NAP: The Network Analysis Profiler, a web tool for easier topological analysis and comparison of medium-scale biological networks. BMC Res. Notes, 10, 278.

13. Gabor Csardi and Tamas Nepusz (2006) The igraph software package for complex network research. InterJournal, Complex Systems, 1695.

14. Pavlopoulos,G.A., Paez-Espino,D., Kyrpides,N.C. and Iliopoulos,I. (2017) Empirical Comparison of Visualization Tools for Larger-Scale Network Analysis. Adv. Bioinforma., 2017, 1278932.

15. Fruchterman,T.M.J. and Reingold,E.M. (1991) Graph drawing by force-directed placement. Softw. Pract. Exp., 21, 1129–1164.

16. Kamada,T. and Kawai,S. (1989) An algorithm for drawing general undirected graphs. Inf. Process. Lett., 31, 7–15.

17. Reingold,E.M. and Tilford,J.S. (1981) Tidier Drawings of Trees. IEEE Trans. Softw. Eng., SE-7, 223–228.

18. Adai,A.T., Date,S.V., Wieland,S. and Marcotte,E.M. (2004) LGL: creating a map of protein function with an algorithm for visualizing very large biological networks. J. Mol. Biol., 340, 179–190.

19. Gavin,A.-C., Bösche,M., Krause,R., Grandi,P., Marzioch,M., Bauer,A., Schultz,J., Rick,J.M., Michon,A.-M., Cruciat,C.-M., et al. (2002) Functional organization of the yeast proteome by systematic analysis of protein complexes. Nature, 415, 141–147.

20. Gavin,A.-C., Aloy,P., Grandi,P., Krause,R., Boesche,M., Marzioch,M., Rau,C., Jensen,L.J., Bastuck,S., Dümpelfeld,B., et al. (2006) Proteome survey reveals modularity of the yeast cell machinery. Nature, 440, 631–636.

21. Pavlopoulos,G.A., O’Donoghue,S.I., Satagopam,V.P., Soldatos,T.G., Pafilis,E. and Schneider,R. (2008) Arena3D: visualization of biological networks in 3D. BMC Syst. Biol., 2, 104.

22. Secrier,M., Pavlopoulos,G.A., Aerts,J. and Schneider,R. (2012) Arena3D: visualizing time-driven phenotypic differences in biological systems. BMC Bioinformatics, 13, 45.

